# Learning multi-cellular representations of single-cell transcriptomics data enables characterization of patient-level disease states

**DOI:** 10.1101/2024.11.18.624166

**Authors:** Tianyu Liu, Edward De Brouwer, Tony Kuo, Nathaniel Diamant, Alsu Missarova, Hanchen Wang, Minsheng Hao, Tommaso Biancalani, Hector Corrada Bravo, Gabriele Scalia, Aviv Regev, Graham Heimberg

## Abstract

Single-cell RNA-seq (scRNA-seq) has become a prominent tool for studying human biology and disease. The availability of massive scRNA-seq datasets and advanced machine learning techniques has recently driven the development of single-cell foundation models that provide informative and versatile cell representations based on expression profiles. However, to understand disease states, we need to consider entire tissue ecosystems, simultaneously considering many different interacting cells. Here, we tackle this challenge by generating *patient-level* representations derived from multi-cellular expression context measured with scRNA-seq of tissues. We develop PaSCient, a novel model that employs a multi-level representation learning paradigm and provides importance scores at the individual cell and gene levels for fine-grained analysis across multiple cell types and gene programs characteristic of a given disease. We apply PaSCient to learn a disease model across a large-scale scRNA-seq atlas of 24.3 million cells from over 5,000 patients. Comprehensive and rigorous benchmarking demonstrates the superiority of PaSCient in disease classification and its multiple downstream applications, including dimensionality reduction, gene/cell type prioritization, and patient subgroup discovery.

## 1 Introduction

Technological innovations in the past decade have led to the collection of vast and exponentially growing amounts of data for biological research, which can help revolutionize our understanding of human disease biology [9,3,40,36]. In particular, the advent of single-cell RNA-seq (scRNA-seq) has enabled the charting of the heterogeneity of cell states and functions, by profiling the expression of hundreds of millions of cells [45]. The large number of cell profiles within and across experiments has opened the way to discoveries from new cell types [22], distinct genes programs associated with response to therapy or drug resistance, specific marker genes [44,34], and unique patient subsets [46,53]. Nevertheless, most scRNA-seq studies were analyzed in isolation and only from a limited number of patients, hindering our ability to understand biological processes at a patient level [28,49,18,19,10,1]. Moreover, studies have typically focused on partitioning cells into categories (types, subtypes, states, etc) and then studying each of them separately, with only limited efforts focused on the overall ecosystem of cells assembled together. Yet, diseases typically involve breakdown of homeostasis in tissue, impacting multiple cells.

Fortunately, the growing number of scRNA-seq studies has now reached a total number of patients that can realistically support machine learning approaches capable of modeling disease biology at a patient level [4,43]. Reasoning about the disease process at the patient level with the granularity of single-cell expression could potentially help uncover subgroups within patient populations (endotypes), understand or predict patient responses to therapies, and advance toward more precise and personalized medicine.

These considerations have motivated the development of machine learning models to aggregate cells to identify disease states. However, existing models only focus on binary disease classification, and were trained with only few samples and studies [15,35,59,38], failing to leverage the large repositories of singlecell expression data available. A more recent work incorporates a larger patient corpus but focuses on multi-modal biomedical data integration, and limits its disease prediction to COVID-19 only [33]. By contrast, we aspire to a method that can leverage the full scope of available data and jointly model all diseases in a single model. However, this vision comes with significant challenges, such as the inherent confounding and batch effects of pooling together data from different studies [30], the imbalanced composition of different tissues, cell types, and diseases [12], and the noise of scRNA-seq data [20,7].

Here, we propose PaSCient, a foundation model that produces a patient representation based on the gene expression of all cells in a patient’s sample, by leveraging large scale single-cell expression studies across different tissues and disease. Intuitively, each patient is represented as a set (or bag) of cells, which our model processes to provide a biologically informed vector representation of the patient. To achieve patient-level representations, we rely on a dedicated attention-based aggregation mechanism and data resampling strategy, which addresses the data integration challenges [55,5] posed by the dataset heterogeneity. Our versatile representation can then be used to compare, cluster, or classify PaSCient: patient representations from single-cell transcriptomics patients. To elucidate disease mechanisms at the patient level, we propose an interpretable mechanism based on integrated gradients [52] to score individual genes and/or cell types in a given patient prediction. This enables a remarkably fine-grained gene or cell-type prioritization, supporting biological discovery at the patient level in terms of individual genes, specific cell types, multiple cell types (simultaneously) and their interconnections. Our comprehensive and rigorous benchmarking further demonstrates the superiority of PaSCient in disease classification compared to single-cell foundation models and underscores its multiple downstream applications, including dimensionality reduction, biological prioritization, and patient subgroup discovery.

To summarize, our contributions are:

1. We propose a machine-learning model that creates patient-level representations based on their single-cell expression profiles. This representation can be used to compare, cluster, or classify patients. Our model leverages single-cell expression studies from over 5,000 patients.
2. The predictions of PaSCient can be interpreted to enable fine-grained prioritization of genes, cell-types, and sets of cell types (and their genes), thereby holistically interrogating disease mechanisms at the patient level.
3. We demonstrate the capabilities of PaSCient on a COVID-19 case study,

showing that the model can be used to infer disease severity subgroups and prioritize cell-type specific genes associated with the disease. Our code is available at https://github.com/genentech/pascient.

## 2 Results

### 2.1 Overview of PaSCient

PaSCient takes the expression profiles of individual cells present within a patient’s sample as input and produces a summarized vector representation of the patient. This representation can then be used for downstream tasks such as dimensionality reduction and visualization, biological feature prioritization, treatment response prediction, and disease severity prediction, among others (Figure 1(a)).

**Fig. 1.**
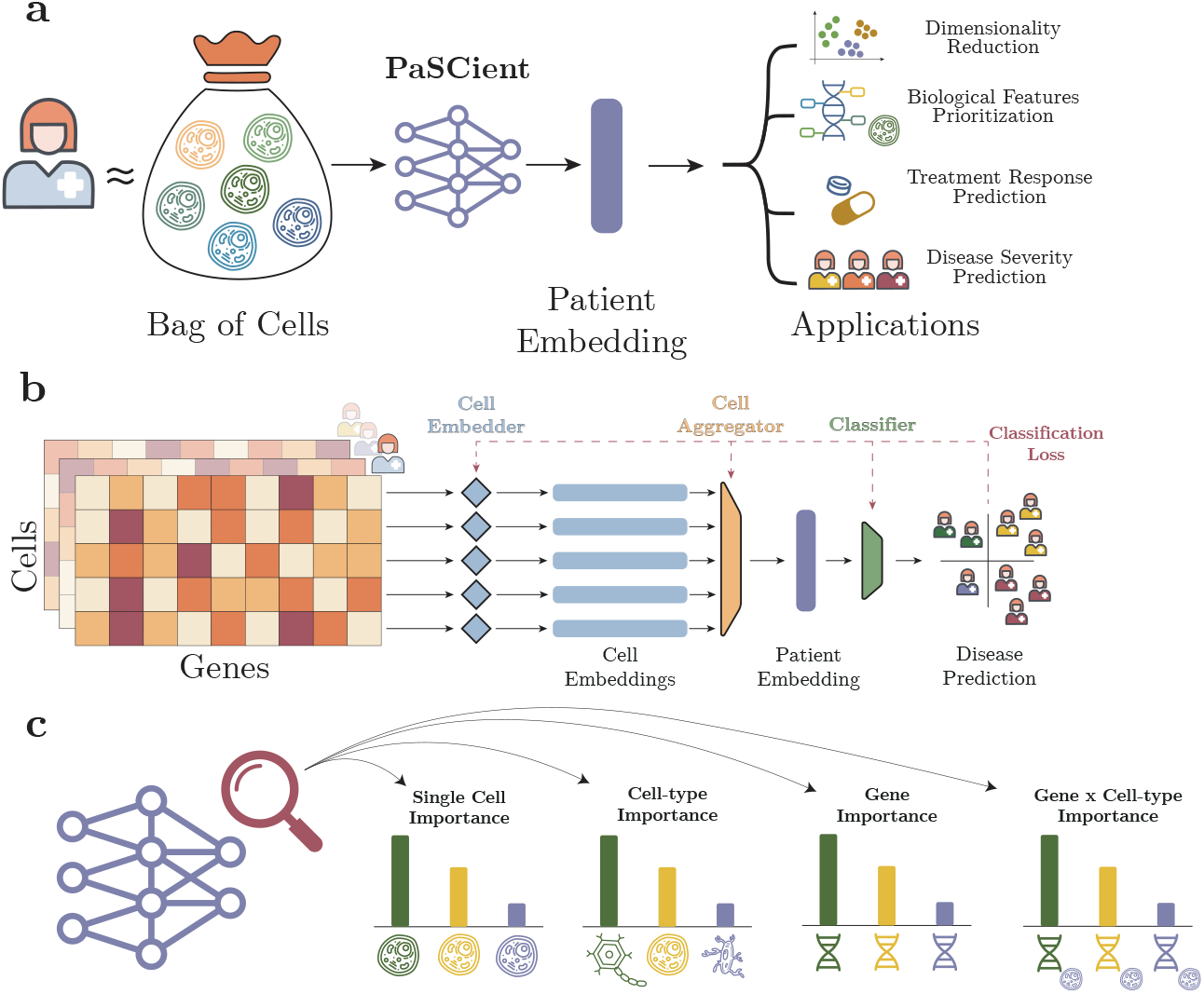
The landscape of PaSCient. (**a**) Model description and applications. PaSCient abstracts each patient as a bag of cells and outputs a single vector summarizing the patient’s cellular context. This vector can be used for various downstream tasks, such as dimensionality reduction, visualization, biological feature prioritization, and predicting treatment response or disease severity. (**b**) Model architecture and training. Each bag of cells is represented as a gene-expression matrix, where rows correspond to individual cells, and columns represent specific genes. PaSCient first embeds each cell individually, and these cell embeddings are then summarized into a patient-level representation by a weighting the embeddings with cell-level attention. A final classifier takes this patient embedding as input to predict the disease status. The entire architecture is trained end-to-end. (**c**) Model interpretability. PaSCient enables finegrained interpretability, generating importance scores at various levels—for individual cells, groups of cells (e.g., cell types), individual genes, or genes within specific cell groups—providing detailed insights into each patient’s cellular landscape.

#### Architecture

The architecture of PaSCient is inspired by DeepSet [63]. The gene expression of the different cells of a given patient *i* is represented as a matrix 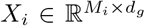, where *M*_*i*_ is the number of cells for patient *i*, and *d*_*g*_ is the number of genes measured. We first encode each cell in the sample using a learnable cell embedder function *f*_*θ*_ : ℝ^*d*^ *d*_*h*_, where *d*_*h*_ is the dimension of the cell representations. At this stage, a patient is represented as a set of vectors {**z**_*j*_ : *j* = 1, …, *M*_*i*_} of size *d*_*h*_. This set can be abstracted as a matrix 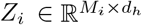. To create a patient-level embedding **e**_*i*_, we used a softmax-attention pooling layer:

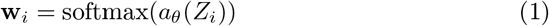

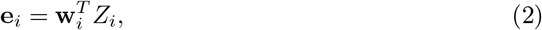

where 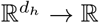 is a neural network acting on each row of *Z*_*i*_ independently. Lastly, the patient-level embedding is fed into a neural network classifier 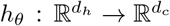, where *d*_*c*_ is the number of disease classes in the pooled dataset. The final disease prediction is obtained as:

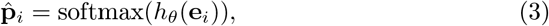

where 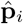 represents the predicted probabilities for each disease label. We train PaSCient end-to-end by minimizing the cross-entropy between predicted diseasestate label and observed disease-state label. Different aggregation mechanisms were investigated during the development of the method. A softmax-attention layer was found to be the most effective in our ablation studies, as shown in Figure 2(b). To address the disease and tissue heterogeneity of the dataset, we introduce a dedicated sampling strategy that gives more importance to sample with low prevalence diseases and tissues. More details can be found in the Methods section.

**Fig. 2.**
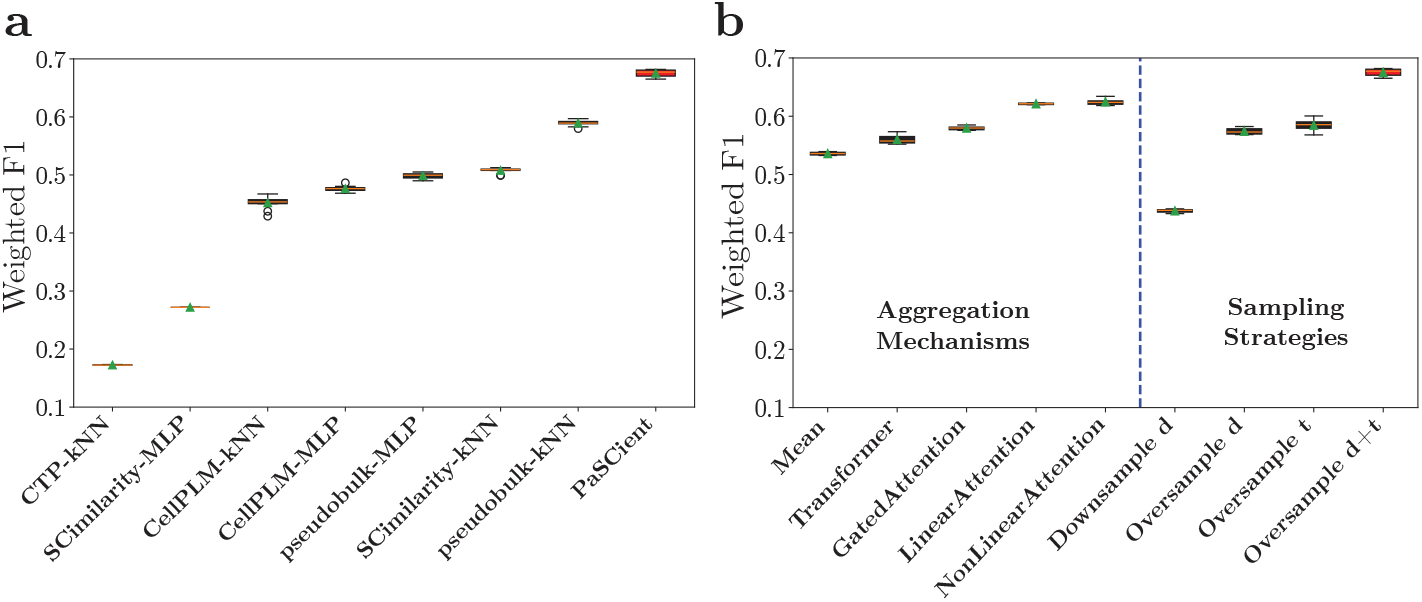
Benchmarking the performance of PaSCient on multi-disease classification. (**a**) Weighted F1-score results. Performance comparison between PaSCient and relevant baseline models, with standard deviations calculated from experiments using different seeds. PaSCientemploys non-linear attention aggregation combined with oversampling based on disease and tissue. (**b**) Ablation studies. Analysis of different training configurations for PaSCient, including various cell-level aggregation methods (without resampling) and sampling strategies to address label imbalance. The best performance was achieved using non-linear attention aggregation with oversampling based on both disease and tissue labels (Oversample d+t).

#### Fine-grained importance scores

To interpret the predictions of PaSCient, we develop an approach relying on integrated gradients (IG) [52]. This procedure starts by producing a gradient attribution for each cell-gene combination of the input sample using IG. Given the resulting matrix of attributions, we average attributions based on different dimensions, leading to different levels of interpretability. For instance, averaging the attributions over genes leads to importance scores for each individual cell, whether averaging over cells leads to importance scores for individual genes. A similar rationale can be employed to generate importance score for groups of cells (or cell types) and individual genes within a given group of cells (Figure 1(c)).

#### Dataset

Our dataset includes 24.3 million scRNA-seq count profiles from over 5,000 patient samples spanning 135 unique disease-state labels, across 413 studies, and 189 tissues (organs). Each patient contributed to a single sample (such that patient and samples can be used interchangeably in this text). All datasets are publicly accessible on CELLxGENE [43]. Cells were all profiled using droplet based scRNA-seq from 10X Genomics. The data were split into a training (60%), validation (20%), and test set (20%), ensuring that all samples from a given study are in the same split. A visual summary of our splits is described in Appendix E. The data distribution was imbalanced in terms of diseases and tissues, *e*.*g*. COVID-19 patients accounted for ∼9% of the samples, while multiple sclerosis only for ∼2% (Extended Data Fig. 2 (a) and (b)).

### 2.2 PaSCient can accurately classify disease from a patient’s scRNA-seq profiles

We train PaSCient to predict the disease label associated with each sample in the dataset and evaluate its performance in terms of weighted F1-score, a widely used metric for evaluating classification performance [2,13]. We compare our approach with different embedding baselines, such as a simple pseudo-bulk approach, using cell-type proportions (CTP), as well as state-of-the-art single-cell foundation models (CellPLM [56] and SCimilarity [16]). For each of these methods, we consider two classifiers to predict the label from the patient embedding: k-Nearest Neighbor Classifier (kNN) [42] and a multi-layer perceptron (MLP).

Remarkably, PaSCient outperforms all baselines by a significant margin (Figure 2 (a)). Notably, a simple pseudo-bulk approach outperforms more complicated foundation models in this task. Additional results on a simpler binary classification task (*i*.*e*., COVID-19 vs. healthy) are given in Appendix F, including the comparison with the most recent domain-expert model ScRAT [35], which performs significantly worse than PaSCient.

We investigated different aggregation mechanisms for pooling cell-level embeddings into a patient-level embedding, including mean-pooling, transformer, gated attention, linear attention, and non-linear attention mechanisms. We found that non-linear attention performed best, improving the weighted F1-score by 16.6% compared to a mean-pooling mechanism (Figure 2(b)). The transformer approach, although more expressive, results in poor performance, probably due to a larger than necessary number of parameters for this task.

To account for the class imbalance in the data, we investigated different resampling mechanisms. We studied the impact of resampling both per disease-class and per tissue-class (Methods). Oversampling the training set for both disease and tissue resulted in a significant improvement compared to baseline (Figure 2(b)). Model training and hyper-parameter tuning details are given in Appendix G.

The patient embedding space learned by PaSCient is organized by disease state (Figure 3(a)) and by tissue (Figure 3(b)). Notably, COVID-19 patients partition into two clusters, corresponding to blood and lung tissue samples. Additional analyses of the patient embedding space, aggregated per disease, are given in Appendices H and I.

**Fig. 3.**
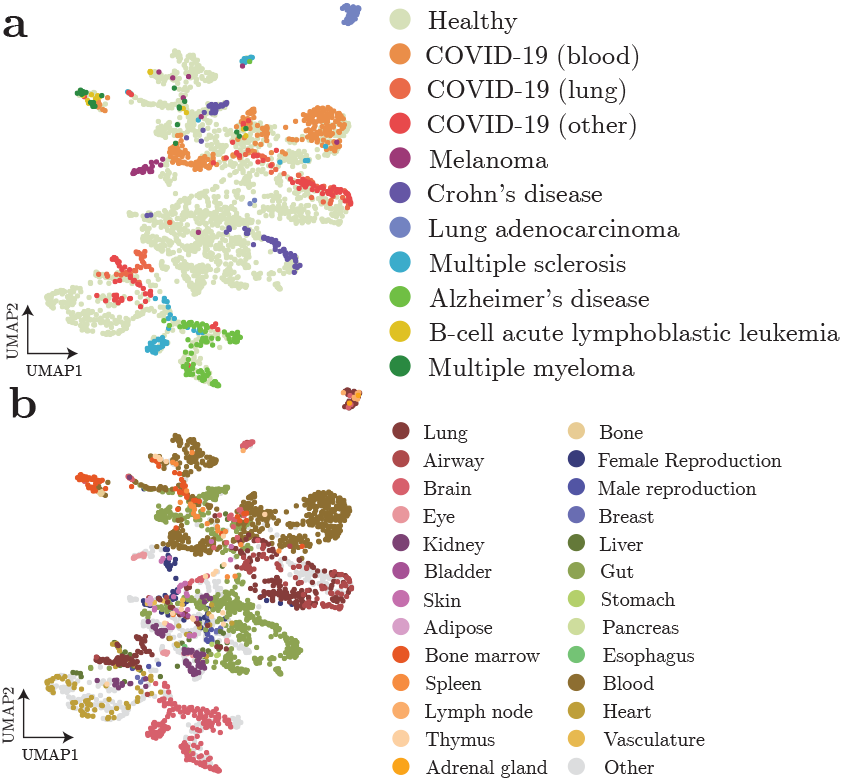
Patient embeddings using PaSCient organize by both tissue and disease. Uniform manifold approximation and projection (UMAP) of patient embeddings colored by each of 8 most common disease labels (**a**) or by tissue (**b**). We only visualize the samples whose disease-state labels exist in all the splits.

### 2.3 PaSCient prioritizes gene and cell-type roles in disease prediction

We use our importance score methodology (described in Section 2.1 and in the Methods section) to enable a fine-grained analysis of the individual cells and genes that contribute most to a disease of interest. As a proof of concept, we focus our analysis on COVID-19 prediction and select a cohort of patients with a COVID-19 disease label.

We first compute cell type level attributions to uncover what cell types were contributing most to the COVID-19 label for each patient (Figure 4(a)). The highest average attributions (computed over all patients) are found for classical monocytes and platelets, suggesting the importance of these cell types in COVID-19. Notably, these cell types have been identified in the literature as playing a key-role in the disease pathogenesis [23,58].

**Fig. 4.**
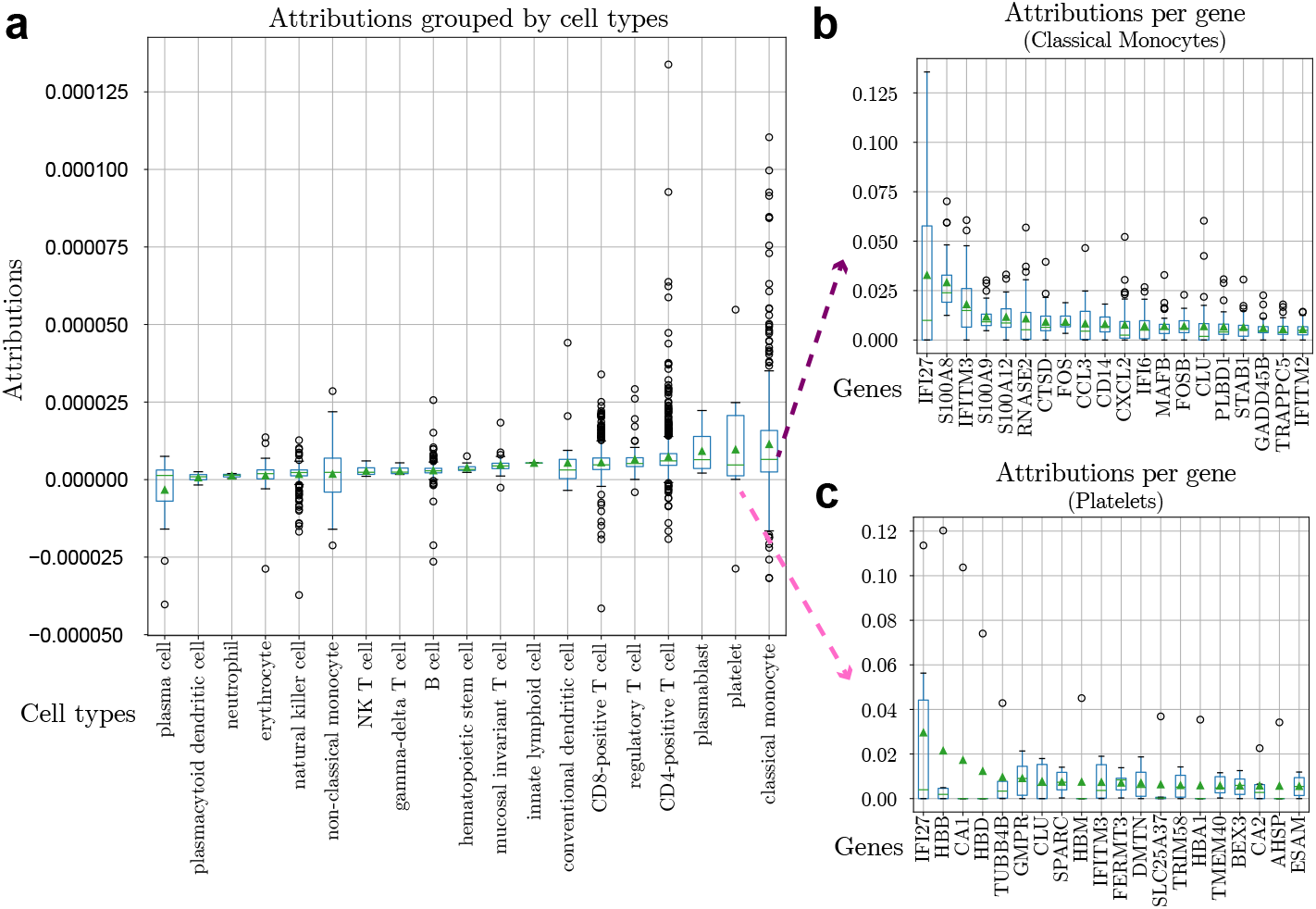
Prioritizing cell types and cell-type-specific genes for COVID-19 by integrated gradients (IG) analysis. (**a**) Attributions averaged over cells and genes for each cell type (each point is a patient). Cell types are ranked by their mean attribution, with classical monocytes and platelets identified as the most predictive for COVID-19 diagnosis. (**b**) Attributions aggregated over classical monocytes (each point is a patient). Genes are ranked by mean attribution, with the green line indicating the median value and the green triangle denoting the mean value. (**c**) Attributions aggregated over platelets (each point is a patient). Genes are ranked by mean attribution, with the green line indicating the median value and the green triangle denoting the mean value.

Remarkably, our fine-grained importance methodology enables further exploration within cell types of interest. We investigate what genes were most impacting COVID-19 prediction for each of these cell types specifically. For each patient, we compute the importance of gene in monocytes (Figure 4(b)) and in platelets (Figure 4(b)). This procedure identifies the specific importance of genes in a given cell type. Ranking genes by average importance reveals that S100A8, IFITM3, and IFI27 are the most pertinent genes in monocytes for COVID-19. IFI27, HBB, and CA1 are found to be most important in platelets. These genes are associated with COVID-19 severity or treatment [37,60,51,64,11].

We validate the set of important genes uncovered by PaSCient by measuring the overlap with the set of differentially expressed genes from ToppCell [21]. A Fisher’s exact test indicates strong overlap for both classical monocytes (p-value=2.1e-22) and platelets (p-value=2.5e-20). A similar analysis for other diseases is presented in Appendix J. These analyses show that we can capture and prioritize disease-specific genes and cell types at different resolutions.

### 2.4 PaSCient recovers disease severity of individual patients

To investigate the patient representations learnt by our method, we collect four scRNA-seq datasets from COVID-19 patients where a severity label is available (mild or severe) [31,50,57,29], and that were not included during training. Visualizing the patient representations generated by our model, we find that the landscape is primarily organized by disease severity and not by study (Figure 5(a)). Conversely, a principal components analysis (PCA) representation of pseudo-bulk data is organized primarily by study rather than severity, highlighting batch effects.

Moreover, the importance scores given by our model to different cell types correlates with disease severity, with significant associations (corrected p*<* 0.01) for NK cells, B cells, myeloid dendritic cells, and MAIT cells. Indeed, there is a significant difference in the magnitude of the integrated gradients attributions of the model, averaged over all myeloid dendritic cells in each patient sample, between mild and severe patient groups (Figure 5(b), Bonferroni-corrected p-value=0.001, rank sum test). Similarly, there is a significant association between the disease severity and the magnitude of the probability of COVID-19 diagnosis predicted by PaSCient (Figure 5(c)). Together, these results show that PaSCient can implicitly represent the disease severity of each patient. Associations between severity and other cell types are given in Appendix K. A case study for predicting drug response is presented in Appendix L.

**Fig. 5.**
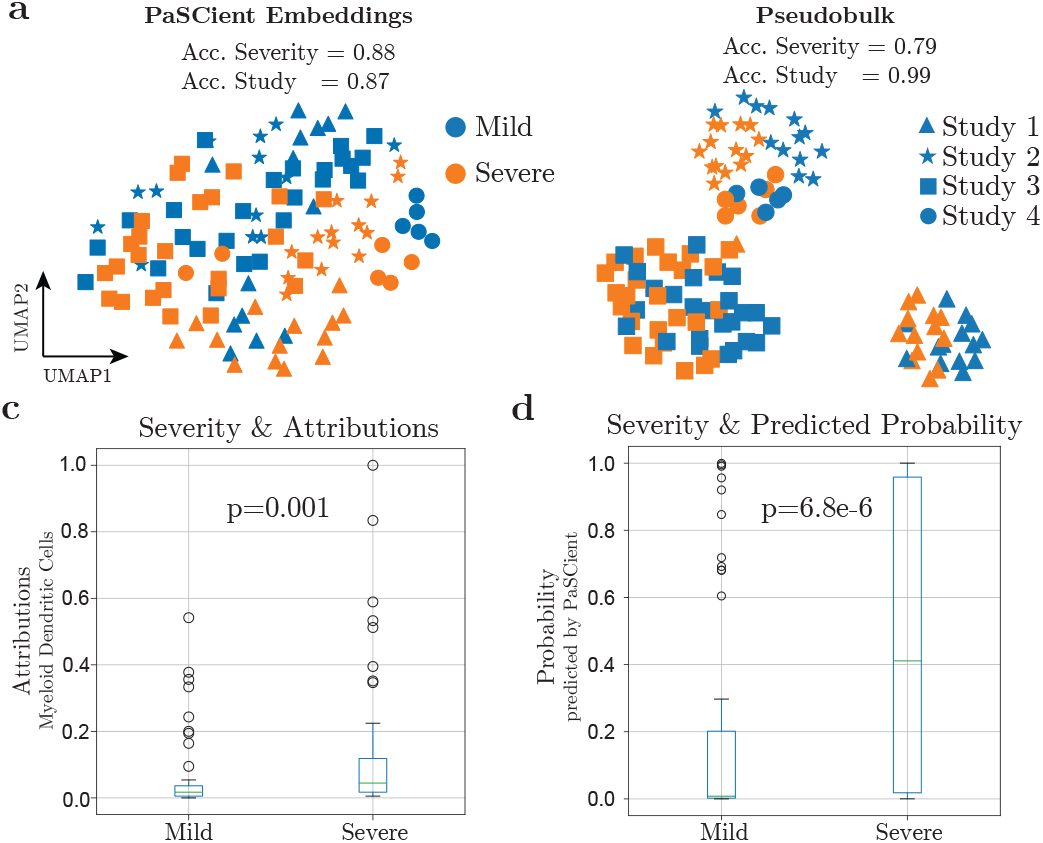
PaSCient captures disease severity in COVID-19 patients. (**a**) Patient embeddings generated by PaSCient and PCA on pseudo-bulk data, colored by disease severity. PaSCient organizes patient representations based on disease severity, whereas the pseudo-bulk embeddings are influenced by study-specific effects. The accuracy of a k-nearest-neighbor (kNN) classifier is reported for both disease severity and study labels to quantitatively assess embedding quality. Higher accuracy suggests the embedding is more organized according to that specific variable. (**b**) Magnitude of integrated gradients attributions averaged across all myeloid dendritic cells for each sample, grouped by disease severity. P-values are Bonferroni-corrected. (**c**) Probability of COVID-19 diagnosis predicted by PaSCient for each sample, stratified by disease severity.

## 3 Discussion

Here, we introduced a new model, PaSCient, that generates patient-level embeddings given a single-cell RNA-seq context, leveraging thousands of samples.

PaSCient builds upon recent single-cell foundation models [8,16,14], and multicellular representations models [15,35,59] but differs in key aspects. First, PaSCient builds upon the large scale training of single-cells foundation models but extends the approach to multi-cellular representations. While single-cell representations can be pooled into a patient-level representation (*e*.*g*., via averagepooling), our experiments showed that this resulted in sub-optimal performance. Our approach is indeed more expressive as it learns a dedicated aggregation mechanism that better reflects the underlying biological processes. Second, PaSCient extends previous works on multi-cellular representations by going beyond binary classifications and by leveraging hundreds of single-cell expression studies. Providing biologically informed patient-level representations presents several advantages for biological and clinical research. Such representations enable a patient-specific understanding of disease mechanisms and can improve patient segmentation, thereby contributing to more targeted therapies. We demonstrated the potential of PaSCient in patient segmentation by showing that the learnt embeddings implicitly encoded clinical information such as disease severity. From a target discovery perspective, we highlighted the fine-grained resolution of our importance scores. We showed that our model could be used to prioritize individual cells and genes, but also groups of cells (such as cell types) and cell-type-specific genes, underlining a promising knowledge discovery toolkit.

Our work represents an important step toward patient-level representations contextualized by single-cell expression. While our datasets included millions of cells, the increasing scale of available single-cell repositories suggests further iterations of this class of models will lead to better representations.

### 4 Methods

#### Notations

We define the aggregated dataset includes *N* patient samples: = {*s*_1_, *s*_2_, …, *s*_*N*_}*”*, where *s*_*i*_ represents the *i*_*th*_ patient sample. Each patient includes *M*_*i*_ cells (where *M*_*i*_ varies per patient): 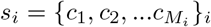, where *c*_*j*_ represents the *j*_*th*_ cell in *s*_*i*_. Lastly, each cell *c*_*j*_ is a vector whose features are gene expression counts with dimension *d*_*g*_ = 28, 231. Each patient can then be represented as a matrix 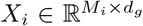. Patient-level metadata is also available such as disease label *y*_*i*_ and tissue label *t*_*i*_.

#### Model architecture

PaSCient combines a cell encoder *f*_*θ*_(·), an aggregator *h*_*θ*_(·), and a classifier *g*_*θ*_(·), all implemented by neural networks. At a high level, the cell encoder produces an embedding for each cell in a patient sample, the aggregator combines the cell embeddings into a patient embedding, and the classifier predicts the disease label based on the patient embedding.

The cell encoder is a linear layer. The classifier is a multi-layer perceptron (MLP) with a final softmax activation. We write the output of the cell encoder as *z*_*i*_ = *f*_*θ*_(*c*_*i*_), the output of the aggregator as **e**_*i*_ = *g*_*θ*_(**z**_*i*_) with 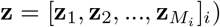) and the output of the classifier as 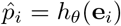. The model is trained by minimizing the cross-entropy between 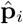 and *y*_*i*_. A graphical depiction of the model architecture is given in Figure 1.

#### Aggregators

We considered multiple different aggregators. Most aggregators have the form of a weighted sum: 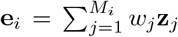. Aggregators differ by the way the weights 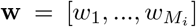 are computed. The mean aggregator uses 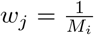; the linear attention aggregator uses **w** = *Softmax*(**z**); the non-linear attention uses *Softmax*(*a*_*θ*_(**z**)) with *a*_*θ*_ a learnable neural network that operates on each **z**_*j*_ independently; and the gated-attention uses **w** = *Softmax*(*U*_*θ*_(**z**) ⊙ *Sigmoid*(*V*_*θ*_(**z**))) with two learnable neural networks *u*_*θ*_ and *v*_*θ*_. The transformer aggregator differs in its architecture as it updates the embeddings of each cell according to the entire sample and sums the resulting embeddings.

#### Resampling strategies

We used the following resampling strategies for addressing the disease and tissue imbalances in the dataset: (1) Downsampling disease: subsampling the most frequent disease classes such as to balance the disease label overall; (2) Oversampling disease: oversampling the least frequent disease classes such as to balance the disease label overall; (3) Oversampling tissue: oversampling the least frequent tissue classes such as to balance the tissue label overall; (4)Oversampling disease and tissue: oversampling the least frequent tissue and disease classes such as to balance both tissue and disease labels overall.

#### Model explainability

We used the integrated gradients method on the input matrix *X*_*i*_ [52]. Computing the integrated gradients on this input results in an attribution matrix 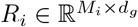 with the same dimensions as the input matrix. The attribution of a given gene was obtained by averaging *R*_*i*_ across all cells. The attribution of a given cell was obtained by averaging over all genes. Any other combination follows from generalizing this procedure.

#### Disease classification metrics

We evaluated classification performance using the weighted F1-score. F1-score is robust to class imbalance and reflects both precision and recall across all classes. Each experiment was repeated 10 times using different seeds leading to different cells being sampled for each patient. This repetition allowed computing an empirical standard deviation on the results.

#### Dataset pre-processing

All datasets were profiled by droplet based scRNA-Seq from 10X Genomics. We removed cell profiles with no gene expression levels and normalized all remaining profiles to the corrected sequencing depth, followed by a *log*(*x* + 1) transformation.

#### Reproducibility and Data

The sources of datasets used for training/validating/testing as well as downstream applications can be found in the Supplementary File 1. Our collected descriptions for diseases and tissues can be found in Supplementary File 2. The genes from ToppCell are listed in Supplementary File 3. The running time of our method for different tasks is included in Supplementary File 4.

