## supplementary for "Learning multi-cellular representations of single-cell transcriptomics data enables characterization of patient-level disease states"

### A Acknowledgements

We thank Jenna Collier, Max Gold, Velina Kozareva, Gokcen Eraslan, Chainglin Wan, David Garfield, and Runming Wei for their suggestions that strengthened the quality of experiments and manuscript.

### B Contributions

TL conceived of the method with input from GH and AR. TL proposed the method and finished experiments with the help of TK, EDB, GS, ND, HW, MH, AM, and HCB. TL and EDB wrote the manuscript with input from GH, GS, AM, and AR. GS, AR, and GH jointly supervised this work.

### C Competing interests

All authors are employees of Genentech or Roche. A.R. is a co-founder and equity holder of Celsius Therapeutics, an equity holder in Immunitas, and until July 31, 2020 was an S.A.B. member of Thermo Fisher Scientific, Syros Pharmaceuticals, Neogene Therapeutics and Asimov.

### D Reproducibility, data, and code availability

The sources of datasets used for training/validating/testing as well as downstream applications can be found in the Supplementary File 1. Our collected descriptions for diseases and tissues can be found in Supplementary File 2. The genes from ToppCell are listed in Supplementary File 3.

We used a server with eight NVIDIA A100 GPUs and 300 GB maximal RAM to conduct all the experiments. The minimal requirement for training/inference based on our model is one A100 GPU, 80 GB. The running time of our method for different tasks is included in Supplementary File 4.

All the codes used in model training and downstream applications can be found in `https://github.com/edebrouwer/pascient`.

### E Dataset

In Extended Data Figure 1, we present a graphical overview of the dataset used for training our model. Extended Data Figure 1 shows the proportions of diseases and tissues in the dataset.

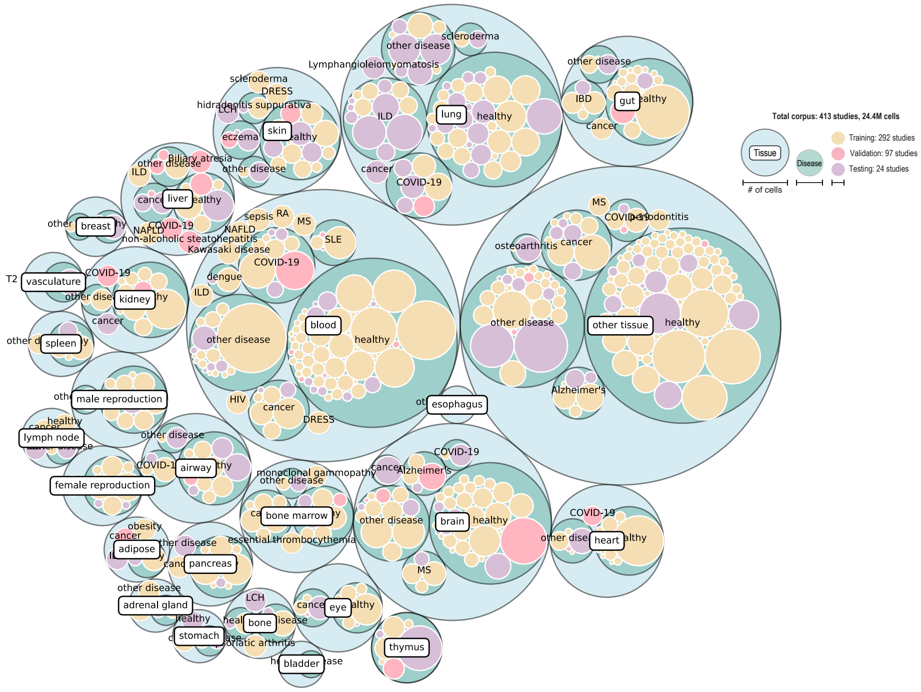


**Extended Data Fig. 1.** Overview of our training datasets colored by tissues and diseases. The length of bubbles represent number of cells included in the given tissue/diseases.

### F Binary disease classification with PaSCient

We evaluated the performance PaSCient to classify patients with health and COVID-19 labels and benchmark it with other methods for performing binary disease-health classification. The results are given in Extended Data Figure 3.

We found that PaSCient outperformed all baselines by a significant margin. We also found that linear attention aggregation performed best than other mechanisms.

#### F.1 Including cell-type information in the prediction

We further investigated whether introducing cell-type information can help predicting the disease label more accurately. We designed a new baseline by using the cell-type proportions to represent each patient and classified the binary conditions based on a kNN classifier (CTP-kNN). We also introduce a modified loss function for PaSCient by using a contrastive learning approach [40] to bring cells from the same type closer together in embedding space, while pushing apart cells of different types (PaSCient-CT). Such an approach can reduce the

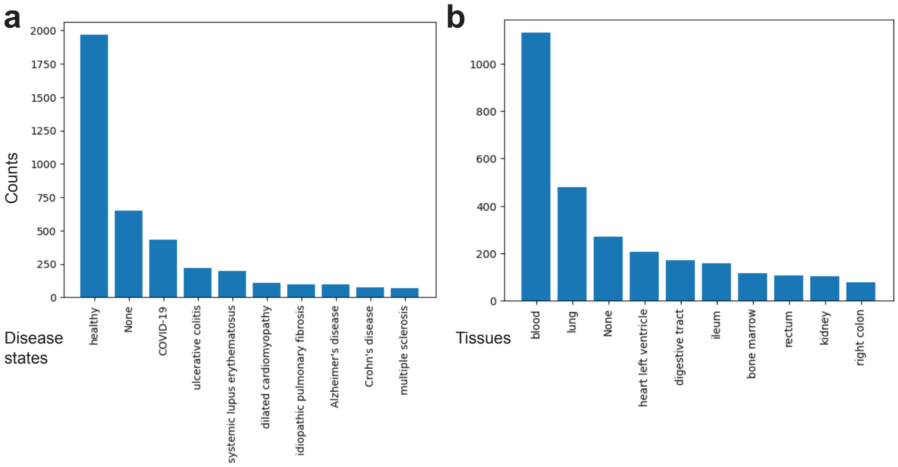


**Extended Data Fig. 2.** Statistics of disease states and tissues in our collected datasets.

potential batch effect existing in the training data [17]. The results are given in Extended Data Figure 3(c). We observed that PaSCient achieved the best performance, followed by our modified loss function (PaSCient-CT) and the cell type proportions baselines.

### G Model training details

During the training process of PaSCient, we explored different factors that could affect the training process, including hyper-parameters, size of sampled cells, composition of diseases, and scaling law [18,28]. The sensitivity analyses focusing on these factors could help us understand the difficulties of patient modeling in a broader vision. We found that learning rate closing to 1e-4 (illustrated in Extended Data Figure 4 (a)) could reduce the negative effects brought by over-fitting. Meanwhile, a small number of epochs ($< 40$ for binary classification and $< 5$ for multi-label classification, based on the epoch with the best validation accuracy) also contributed to better model performances, which matched recent analyses in foundation model training [61]. The dropout rate and weight decay rate also work better with a small value (illustrated in Extended Data Figures 4 (b) and (c)). Meanwhile, we found that increasing the number of sampled cells does not always enhance the performances of PaSCient, shown in Extended Data Figure 4 (d) for the experiments based on binary classification, which also matched previous research about selecting the random sampling policies for multiple instance learning [54]. A suitable range of sampled cell numbers was found to be in $(100, 2000)$.

In the multi-class setting, we faced a more complicated condition with different compositions of diseases with different difficulties. Therefore, to investigate

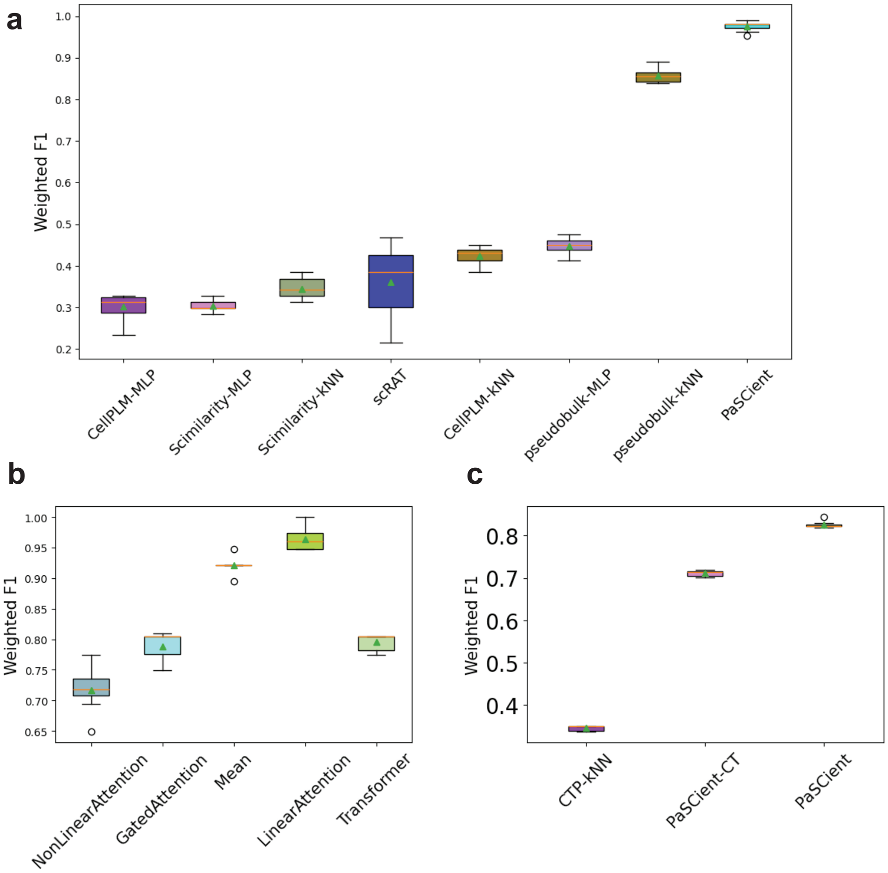


**Extended Data Fig. 3.** Benchmarking results for binary classification. (**a**): Comparisons between PaSCient and the rest of baselines for classifying COVID-19 versus healthy condition. (**b**): Ablation study for different aggregation mechanisms. (**c**) Impact of cell type labels on performance. (PaSCient-CL) uses a modified contrastive training approach that uses cell type labels. CTP-kNN is kNN disease classifier based on cell types proportions for each sample.

the prediction performances specific to different diseases, we hypothesized that model performances was correlated to the frequency of diseases included in the training dataset. PaSCient should be able to classify healthy conditions with diseases whose frequency is low (as multiple binary classification problems). In contrast, PaSCient might face a more and more challenging problem when introducing more diseases in the classification problem (as multiple multi-label classification problems). To validate the first assumption, we separated the eight diseases into eight classification problems versus healthy conditions and trained PaSCient to distinguish them. The results of this experiment are presented in

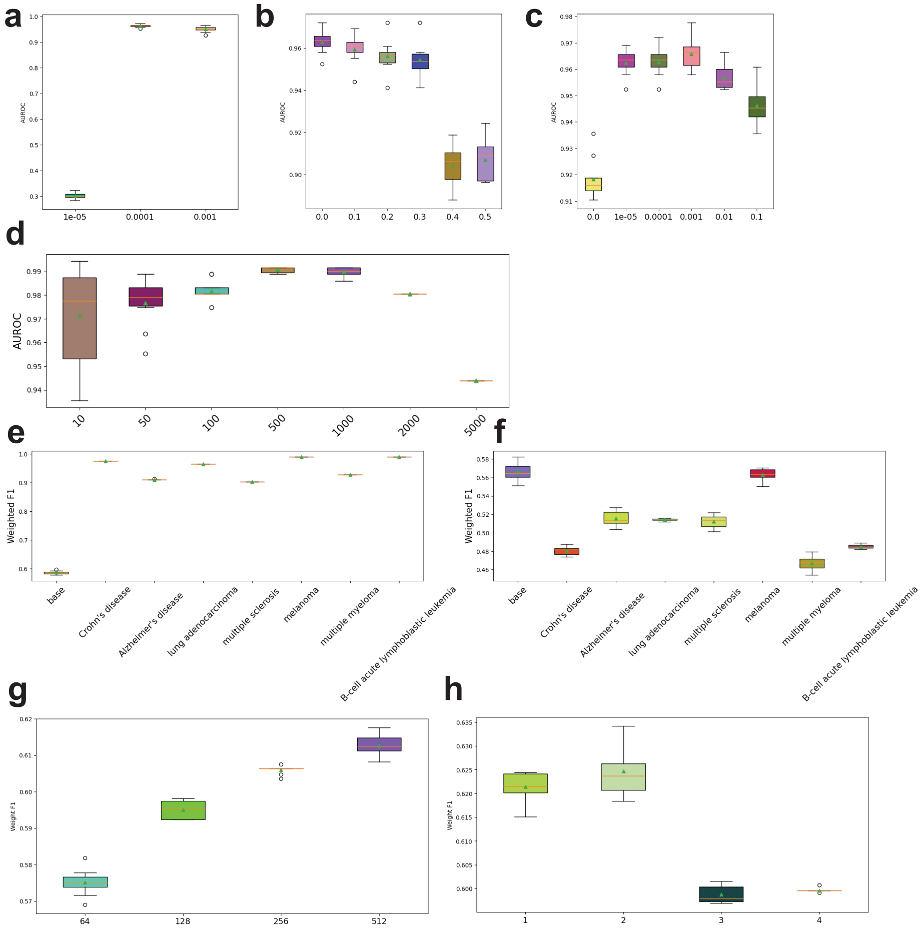


**Extended Data Fig. 4.** Sensitivity analysis of PaSCient. (**a**) Performance of PaSCient under different learning rates for binary classification. (**b**) Performance of PaSCient under different dropout rates for binary classification. (**c**) Performance of PaSCient under different weight decay rights for binary classification. (**d**) Performance of PaSCient under different number of sampled cells for binary classification. (**e**) Performances of PaSCient under the cumulative setting of different diseases states. (**f**) Performance of PaSCient for classifying healthy samples and other samples with different diseases. (**g**) Performance of PaSCient under different widths of neural network layers. (**h**) Performance of PaSCient under different depths of neural network layers.

Extended Data Figure 4 (e). We found that diseases with lower frequency were generally easier to identify. Meanwhile, we added diseases to evaluate the cumulative performances from the largest frequency to the lowest frequency shown in Extended Data Figure 4 (f), which showed that introducing more diseases might reduce classification performances.

To explore the scaling law of PaSCient, we considered adjusting two architecture parameters, including (1) the width of layers (shown in Extended Data Figure 4 (g)) and (2) the number of layers (shown in Extended Data Figure 4 (h)). We found that increasing the width of the layers can help improving the prediction performance, while increasing the number of layers might harm the prediction performance.

### H PaSCient learns the differences and similarities of different diseases in the representation space

Modeling patients is a complex problem, so using classifier performance to measure patient representation is insufficient to show that we have learned meaningful patient embeddings. For the given group of patient embeddings, We can categorize them by disease as well as by tissue of their origin to study the effects of the same disease on different tissues. By averaging the patient embeddings by both diseases and tissues and computing the correlation matrix, we visualized the correlation results in Figure 5. We found that disease embeddings from COVID-19 and lung adenocarcinoma have a high correlation across different tissues, which implied that PaSCient successfully learned the patient sample representations across different tissues from the same disease. Our embeddings also captured signals of different tissues within the same disease demonstrated by the results of hierarchical clustering. Such findings could also be supported by recent research about the multi-tissue damages of COVID-19 [7] and lung adenocarcinoma [65], as these diseases tended to affect different tissues jointly.

### I Evaluating patient embeddings with large language models

Rigorously evaluating the quality of the embeddings produced by a given method is a challenging task, that requires meta-data annotations that is not always available. To address this challenge, we constructed a disease similarity measure based on the text descriptions of each disease, extracted from the Kyoto Encyclopedia of Genes and Genomes (KEGG) database [27,25,26] and National Center for Biotechnology Information (NCBI) [48]. Each disease description was converted to an embedding using the OpenAI text-embedding tool [42]. Similarity between diseases was then obtained by computing the Pearson Correlation Coefficients (PCCs) between their respective embedding. The resulting similarity matrix is given in Extended Data Figure 6(a).

We then constructed a similarity measure between diseases from the embeddings of PaSCient by averaging the embbedings of all patients with a given disease and computing the pairwise Pearson Correlation Coefficients. Lastly, we used a similar procedure for computing disease similarities from pseudobulk data. The resulting similarity matrices are presented in Extended Data Figure 6(a).

To quantitatively assess the discrepancy between the text-based disease similarities and PaSCient-based similarities, we computed the PCCs between both

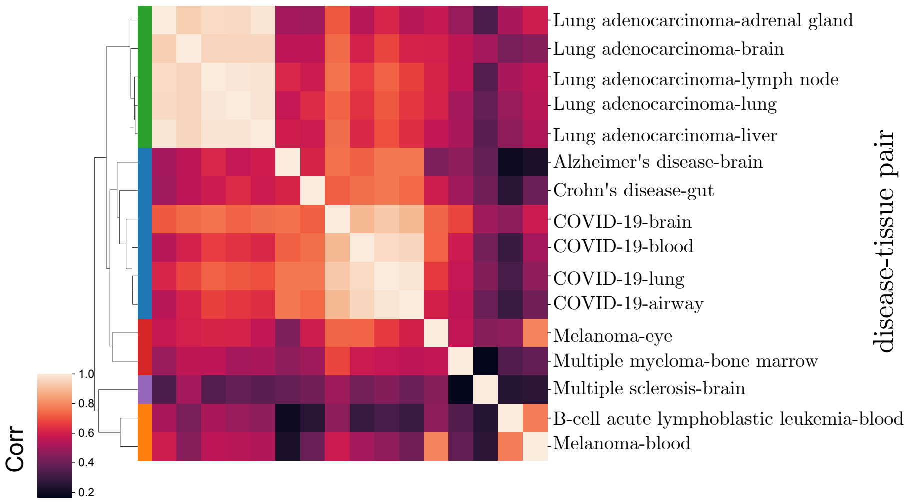


**Extended Data Fig. 5.** Demonstration of patient-sample-level embeddings. Correlation coefficients of disease embeddings across different tissues. The clusters are generated based on the hierarchical clustering with one minus correlation coefficient as distance.

similarity matrices. We found that the correlation from PaSCient's result (PCC=0.65, p-value=9.5e-33) was higher than the correlation from pseudobulk gene expression levels (PCC=0.28, p-value=2.6e-6), suggesting that PaSCient can learn better patient representations by considering both the differences and similarities across different diseases.

Furthermore, we investigated whether PaSCient could capture the similarity of certain disease-tissue embeddings with other embeddings by visualizing the relationship between the correlation from text embeddings and the correlation from PaSCient (Extended Data Figure 6(b,c)). We found that PaSCient aligned with the human understanding of diseases with patient representations from transcriptomic data for the lung samples with lung cancer (PCC=0.83, p-value=1.2e-4) and for the lung samples with COVID-19 (PCC=0.72, p-value=2.0e-3). Overall, these preliminary results show that text embeddings can be a promising metric for evaluating the reliability of learned patient representations.

### J Explainability of multi-disease states

We visualized the importance scores across cell states and genes by different diseases in Extended Data Figure 7 and Extended Data Figure 8.

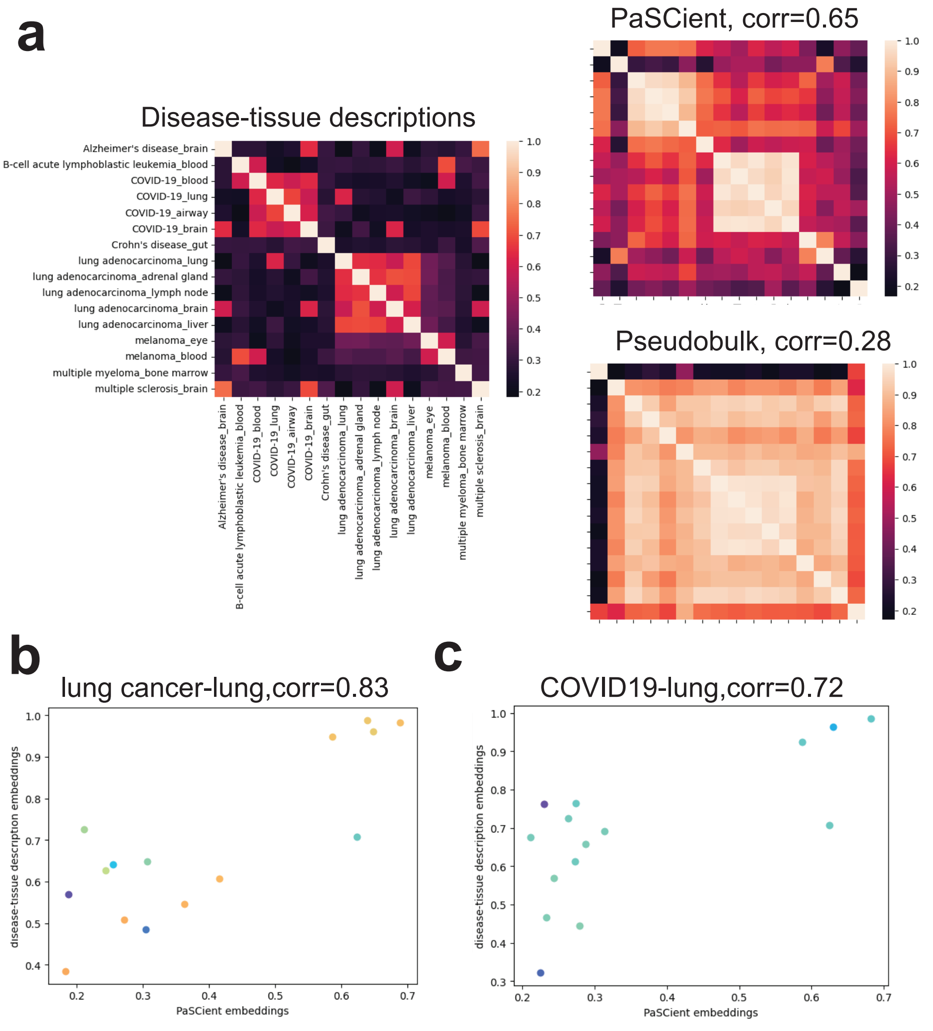


**Extended Data Fig. 6.** Evaluation patient embeddings with the prior information from text descriptions. (**a**): Correlation matrices of embeddings computed based on different rules for disease similarity across different diseases. The sources and correlations between computed matrix and ground truth matrix are labelled in the figure. (**b**) Scatter plot between the correlation computed with embeddings from PaSCient and the correlation computed with text embeddings for describing the similarity of lung cancer in lung and other disease-tissue pairs. (**c**) Scatter plot between the correlation computed with embeddings from PaSCient and the correlation computed with text embeddings for describing the similarity of COVID19 in lung and other disease-tissue pairs. All the p-values corresponding to correlations are significant.

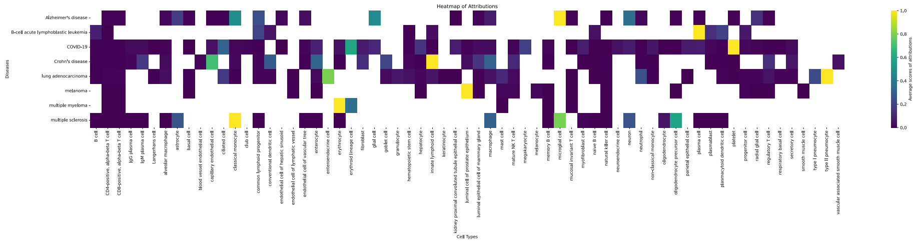


**Extended Data Fig. 7.** The averaged attributions of different cell types for diseases. The attributions are scaled for each diseases for better visualization, and the cell types are corrected and annotated with SCimilarity.

### K Association of cell-types importance scores and COVID-19 severity

We visualized the association between attribution scores and COVID-19 severity for each cell-type in Extended Data Figure 9.

### L Case study: Identification of treatment responders in melanoma

To complement our quality-assessment of the patient representations learnt by PaSCient, we investigated whether it could identify treatment responders for T-cell immunotherapy in melanoma. We collected two scRNA-seq datasets from patients with melanoma treated with T-cell immunotherapy, and for which a binary treatment outcome label was available [47,62].

Following the settings of [33], we utilized the former dataset as a training dataset and the latter as a testing dataset. For reference, we used the pseudobulk data from original expression profiles by patients with a Support Vector Classifier (SVC), and extracted the patient representations by querying PaSCient with the original gene expression profiles and performed classification under the same training/testing datasets. The whole workflow is shown in Extended Data Figure 10 (a). We also visualized the sample embeddings of training dataset in Extended Data Figure 10 (b) and observed clusters for non-response samples. Furthermore, we visualized the classification results in Extended Data Figure 10 (c), and the SVC using representations form PaSCient as input could outperform the baseline model under different classification metrics, especially because the baseline model predicted all patient samples as drug-responsible.

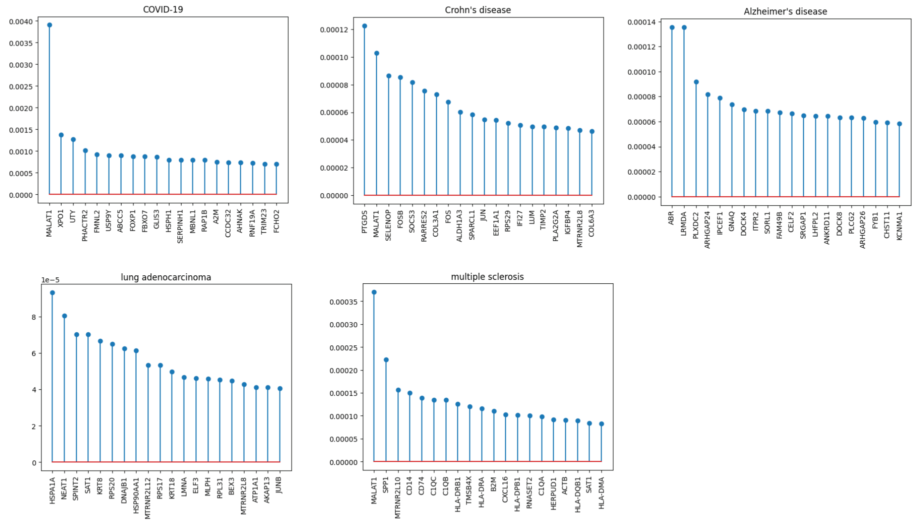


**Extended Data Fig. 8.** Averaged attributions of each gene for different diseases. Each figure represents one disease and we select top 20 genes to present, ranked by their average attributions.

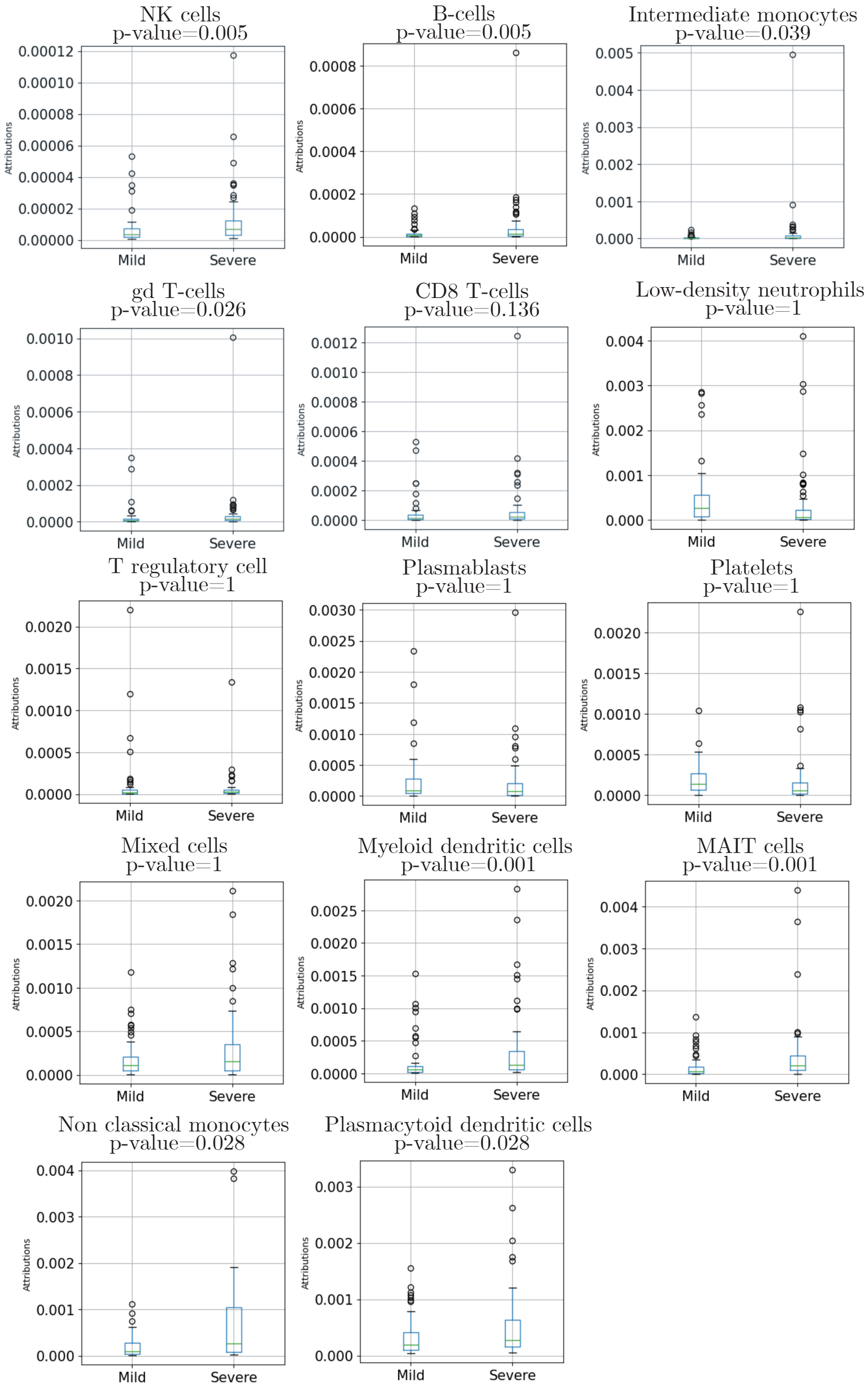


**Extended Data Fig. 9.** Results of t-tests for the association between attribution scores and COVID-19 severity for each cell-type. p-values are Bonferroni corrected.

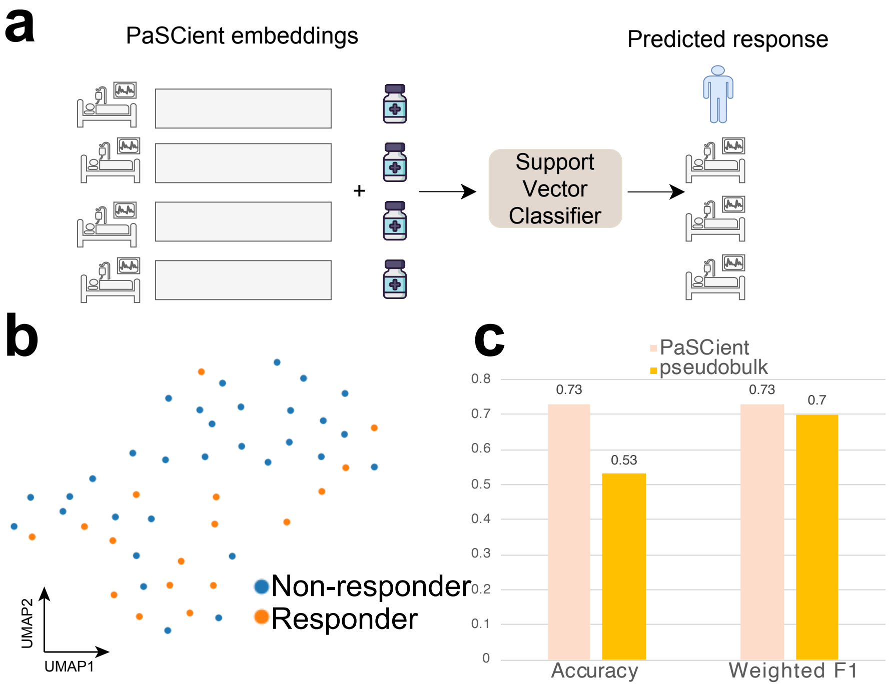


**Extended Data Fig. 10.** Performances of PaSCient for patient-level treatment response prediction. (**a**): Overview of treatment prediction task. We utilize patient sample embeddings from PaSCient as input and classify these samples with their known treatment responses and transfer the knowledge to predict unknown treatment responses in the testing dataset. (**b**): Visualization of sample embeddings colored by treatment responses. (**c**): Benchmarking results between PaSCient embeddings and pseudobulk gene expression levels as inputs for this task.
